# Mapping giant-tree density in the Amazon

**DOI:** 10.1101/2025.02.25.640223

**Authors:** Robson Borges de Lima, Diego Armando S. da Silva, Matheus Henrique Nunes, Paulo R. de Lima Bittencourt, Peter Groenendijk, Cinthia Pereira de Oliveira, Daniela Granato de Souza, Rinaldo L. Caraciolo Ferreira, José A. Aleixo da Silva, Jesús Aguirre-Gutiérrez, Toby Jackson, João R. de Matos Filho, Perseu da Silva Aparício, Joselane P. Gomes da Silva, José Julio de Toledo, Marcelino Carneiro Guedes, Danilo R. Alves de Almeida, Niro Higuchi, Fabien H. Wagner, Jean Pierre Ometto, Eric Bastos Görgens

## Abstract

Tall trees are pivotal in maintaining tropical forests’ biodiversity and regulating biogeochemical cycles, mainly through their influence on carbon storage, nutrient cycling, and habitat structuring. Despite their ecological importance, the drivers behind their spatial distribution and the environmental factors driving the density of these emergent trees remain unexplained. Here we analyzed tall-tree density data derived from airborne LiDAR across 900 randomly distributed transects throughout the Brazilian Amazon. We paired each transect location with climatic, topographic, atmospheric, and soil metrics to investigate regional-level environmental drivers of large tree density using spatial modeling. Water availability and stable climatic conditions (low incidence of storms and of lightning) were the main drivers of tall-tree density at the regional level. The occurrence of tall trees is aggregated along the Amazon, with 14% occurring within 1.3% of the Amazon and 50% concentrated in 11.2% of the area. These patterns suggest that tall trees dominate localized regions of the biome and that high densities of tall trees are primarily dependent on mechanical rather than physiological drivers. Understanding the dynamics of 55.5 million emergent trees is imperative for advancing research in biogeochemistry, ecology, and climate resilience. Our results are especially relevant in the context of conserving nature and carbon stocks under increasing anthropogenic and environmental pressures.

## 1 Introduction

Understanding the density and distribution of tall emergent trees in the Amazon is vital for predicting the carbon balance of Amazonian ecosystems with global environmental change (1–3). Particularly, the local and regional variations in the density of large trees are strongly linked to spatial variations in the above-ground biomass of tropical forests (4, 5) and regulate the microclimate, water availability, light intensity, and understory species diversity (6–10). Around 390 billion trees are estimated to inhabit the Amazon (11), However, large uncertainties remain regarding how many tall emergent trees reach the uppermost stratum of Amazonian forest canopies and how their survival is locally and regionally distributed. These large trees take centuries to reach such sizes, and the species may be unable to regenerate on timescales relevant to the planet’s rapid climate change (12–14). Their longevity, combined with increasing forest degradation and rising temperatures, prevents them from recovering at the same rate they are lost, making them especially vulnerable to current environmental pressures (15). Analyzing the density of large trees is thus fundamental to understanding the dynamics and stability of forest ecosystems.

The density of tall trees is largely influenced by variables that determine their growth and survival rates, encompassing both genetic factors and environmental conditions such as climate, soil, and topography, which shape their structural characteristics (16–19). Moderate temperatures and increased light availability (i.e., providing more energy for growth), alongside low water stress are factors that can favor a large density of tall trees (20, 21). However, the future survival of large trees may be threatened by climate change and land-use changes. For example, large climatic and atmospheric oscillations may lead to physiological and mechanical instability in the structure of tall trees (22, 23). The efficient vascular systems of tall trees are particularly vulnerable to prolonged droughts, which can lead to the collapse of water and nutrient transport (24, 25), and high turbulence caused by strong winds and a high incidence of lightning kill tall trees disproportionately (26, 27). Therefore, the capacity of Amazonian forests to shelter tall trees probably depends on a delicate balance between environmental factors associated with resource availability that promote tree growth to its potential maximum size, alongside mechanisms that promote survival under vegetation disturbances (28). Analyzing the density of tall trees as a function of environmental determinants is crucial to understanding the Amazon’s stability under increasing environmental pressures and climate change.

Field inventories have long been used to map patterns and assess forest diversity in the Amazon, providing critical insights into the composition and distribution of tree species (18, 29–31). However, traditional approaches, such as forest inventory plots do not provide sufficient coverage for evaluating the density of tall trees. This is mainly due to the logistical challenges and precision limitations associated with detecting these trees, which often require large sampling plots, particularly in remote and inaccessible regions (32, 33). While such studies are invaluable for understanding the processes and mechanisms by which trees respond to environmental factors, they often fail to capture tall trees’ full spatial variation and density across the Amazon.

The advent of LiDAR (Light Detection and Ranging) has transformed forest assessments by providing high-resolution, three-dimensional data on forest structure over large spatial scales. This technology can be used for rapid and accurate estimates of canopy height and aboveground biomass in remote regions (34–36). The “Biomass Estimation in the Amazon” (EBA) program, for example, mapped 900 transects, covering 375 hectares each, which represents a remarkable 1,000-fold increase in sampling capacity compared to traditional forest inventory plot methods (Fig. 1). This technological breakthrough has, for the first time in the Amazon, enabled more precise and accurate mapping of biomass stocks (37) and reshaped our understanding of forest structure and the distribution of emergent trees in the Amazon (28, 38). In particular, Gorgens et al. (38) identified the tallest tree ever recorded in the Amazon — a *Dinizia excelsa* reaching 88.5 m height in the eastern portion of the Roraima biogeographical province in Brazil. This discovery underscored the ecological importance of these towering giants and highlighted the critical role remote sensing techniques have in uncovering previously unknown aspects of tropical forest ecosystems. Despite these advances, there is still a dearth of LiDAR-based estimates of large tree density across the tropics. Better estimates of large-tree density is crucial to improve our understanding of the ecological processes and mechanisms driving tree-environment interactions, that are vital for predicting the carbon cycle dynamics and species vulnerability of Amazonian forests.

**Figure 1:**
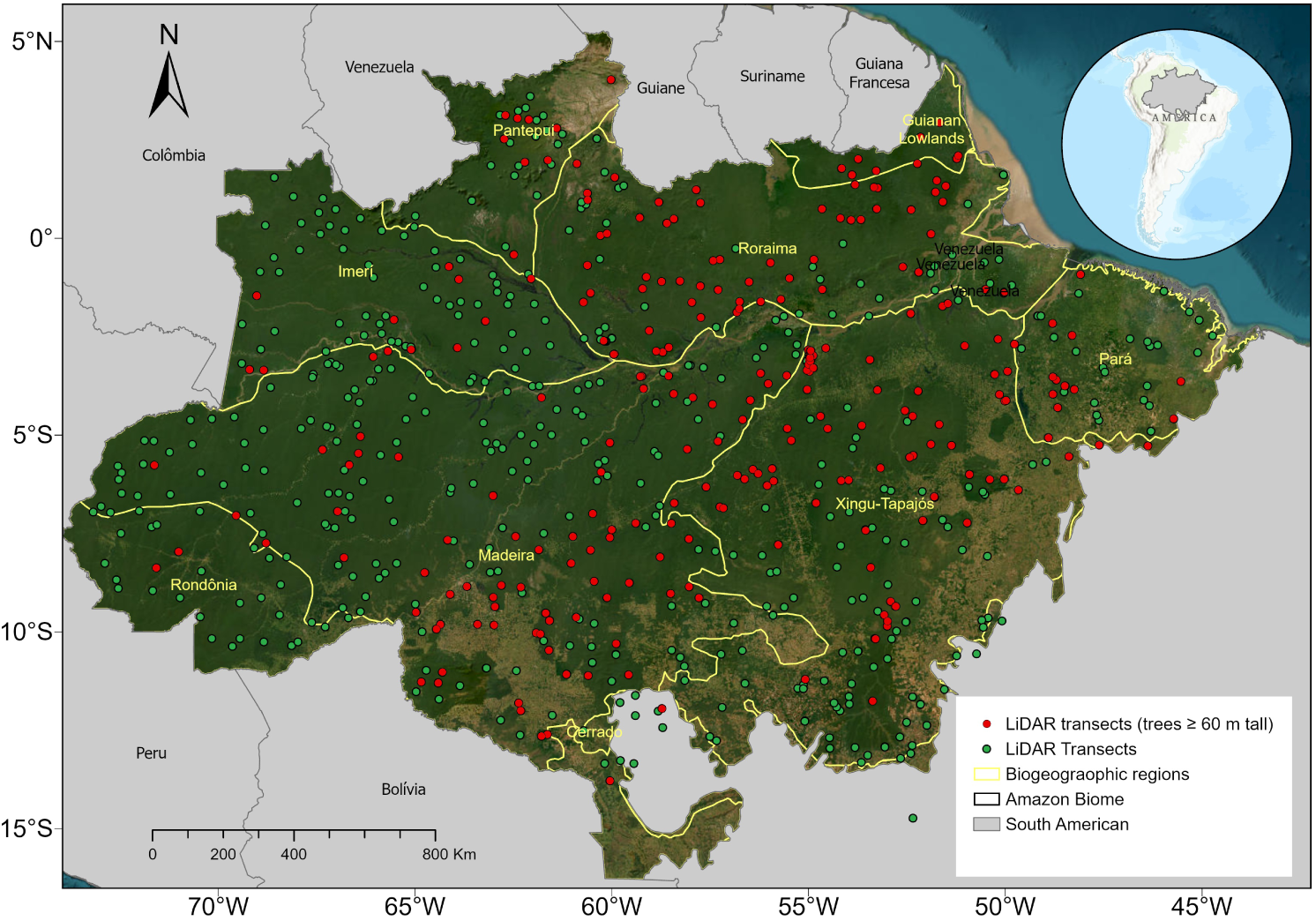
Distribution of transects mapped by airborne LiDAR across the Amazon Basin. The figure shows 900 randomly distributed transects covering the eight biogeographic provinces in the region. Red points represent transects where trees taller than 60 meters were identified, while green points indicate transects without trees ⩾ 60 m. Yellow lines delimit major biogeographic regions, including Imeri, Roraima, Guiana Shield, Pantepui, Madeira, Rondônia, Xingu-Tapajós, and Pará. The LiDAR data transects is available at zenodo.org

Here, we generated the first map of the spatial distribution of the tall tree density (height ⩾ 60) in the Amazon basin, integrating spatially explicit climate, topography, atmosphere and soil information. We first estimate tall-tree density in biogeographic provinces of the Brazilian Amazon using a random forest machine learning method. Then, we assessed the underlying environmental factors and identified the principal components that explain most of the variance in tall-tree density in these biogeographic provinces. We also analyze the broader implications of these results for biomass estimates. These comprehensive analyses provide new insights into how different environmental variables influence tall-tree density and provides crucial information for forest conservation and management policies in the Amazon.

## 2 Results

### 2.1 Density of tall trees and model analysis

LiDAR mapping detected 5,522,736 trees, with an approximate average of 767 trees/km2 in approximately 7,202 km2 of overflown area. The maximum height found was 88.5 meters, validating this information in the field as the tallest known tree in South America (38). The observed density of trees ⩾ 60 per transect ranged from 0 to 227 trees/km2, with an average density of 0,8 trees/km2 *(SI Appendix, Fig. S2a)*

Our analysis confirmed, with a high level of accuracy, that the fitted Random Forest model (k-fold cross-validation: RMSE = 4.81 trees/km2) indicates a general spatial trend in the density of tall trees throughout the Amazon basin. The spatial model explained 79% of the variation in tall tree density and presented a significant correlation between the observed and estimated values *(SI Appendix, Fig. 2b)*, of which the residuals did not show significant spatial autocorrelation *(SI Appendix, Fig. 2c)*. The results from the spatial cross-validation demonstrated satisfactory performance of the Random Forest model in estimating the density of tall trees. The root mean square error (RMSE) was 10.49 trees/km2, indicating reasonable accuracy in the predictions. In contrast, the mean coefficient of determination (R2) of 0.55 shows that the model could explain a substantial portion of the variability in the observed data. Additionally, the mean bias of 0.35 reveals a low systematic deviation, suggesting that the model showed good overall accuracy without tendencies toward underestimation or overestimation.

The spatial model prediction of large tree density shows differences among biogeographic provinces, Fig. 2, with markedly higher values in the northeastern portion of the Amazon, where there is a predominance of terra firme forests in the northern Guiana Shield and the eastern portion of the Roraima biogeographic provinces (darker area on the map, approximately -58*^◦^* to -51*^◦^* longitude, and -4*^◦^* to -3*^◦^* latitude). The density of tall trees in these areas may exceed 120 individuals/km2, estimated by the spatial model (Fig. 2). Note that these regions are marked by low incidence of lightning and lower wind speeds *(SI Appendix, Fig. 4)*. Densities close to zero can be seen in large portions of the south-central eastern Amazon (Pará, Xingu-Tapajós and Madeira provinces; note that these biogeographic provinces coincide with the arc of deforestation). We also noted lower densities throughout the Imerí province and the entire northern portion of the Madeira province. This westernmost strip of the biome is marked by the solid influence of the seasonality of the dry season *(SI Appendix, Fig. 4)*. Given the 4.21 million km2 of the Brazilian Amazon biome, the spatial model estimates approximately 55.5 million trees taller than 60 meters (0.0001% of the estimated ≈ 390 billion trees for the Brazilian Amazon biome), with an average density of 13.5 (± 4.5) tall trees/km2. However, we found a sizeable spatial aggregation of these tall trees, with approximately 14% of tall tree density occurring in 1.28% of the total area of the Amazon (latitudes -2*^◦^* to 3*^◦^* S and 53*^◦^* to 58*^◦^* W) and 50% occurring in 11,2% (latitudes -5*^◦^* to 5*^◦^* S and 51*^◦^* to 60*^◦^* W).

**Figure 2:**
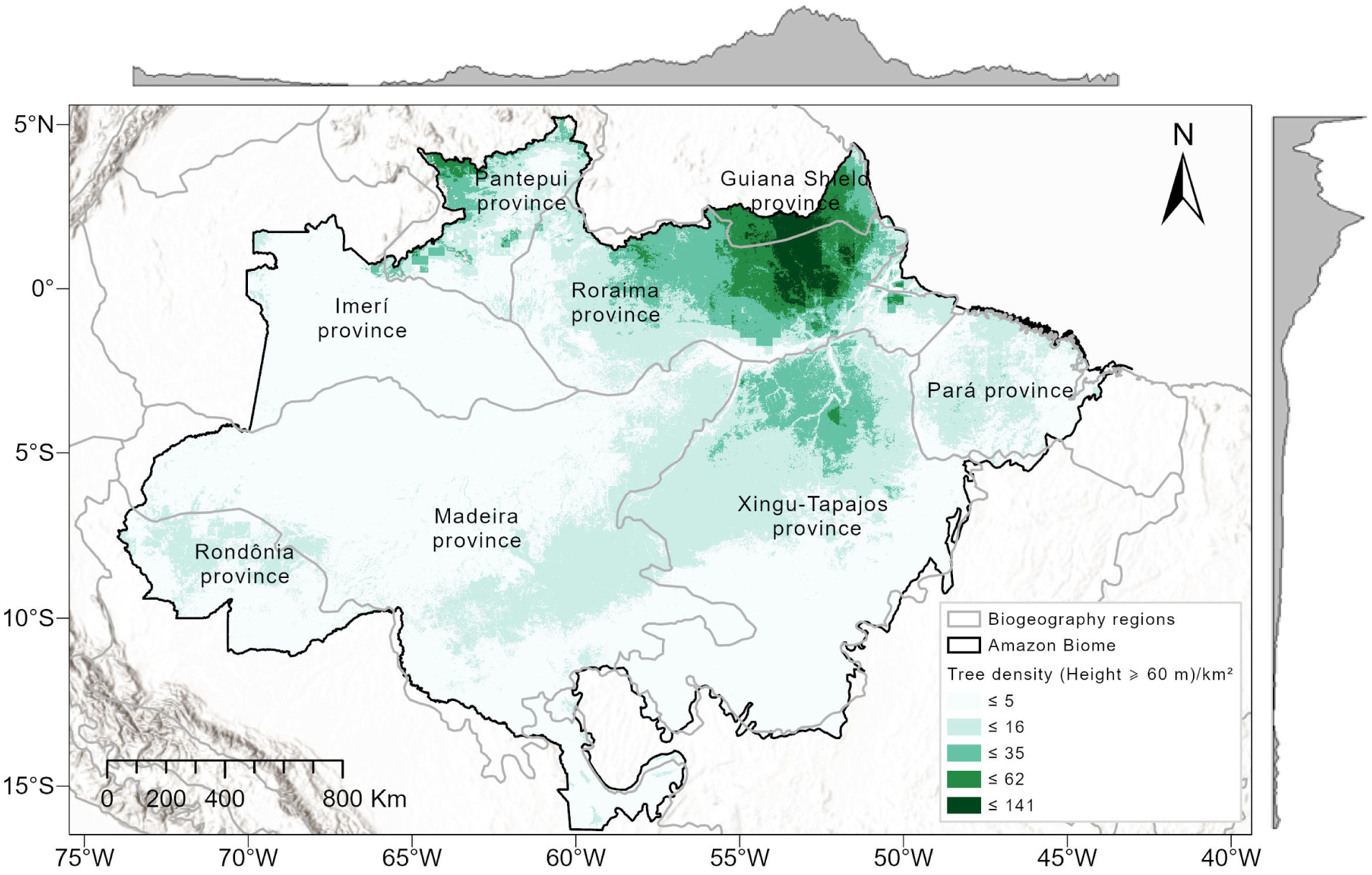
Potential map generated by the random forest spatial model to define ideal zones for the occurrence of high density of giant trees in the Amazon. The areas are color-coded according to different density ranges, where: Lighter shades indicate lower tree density (*<* 5 trees/km2), Darker shades indicate higher density (up to 141 trees/km2). Grey lines delimit major biogeographic provinces within the Amazon biome (such as the Guiana Shield, Xingu-Tapajós, and Roraima), showing distinct density patterns across the Amazon region. The map is available at doi.org/10.5281/zenodo.13850976

The essential variables according to %IncMSE for the model were Mean annual precipitation (mm) (pannual) (13.01%), Mean annual temperature (°C) (tannual) (11.89%), zonal wind speed (W-E, *m s^−^*^1^) (uspeed) (11.65%) and Fraction of clay content (clayContent) (10.98%) *(SI Appendix, Fig 3b)*. Meridional wind speed (N – S, *m s^−^*^1^) (vspeed) and lightning rate (flashes rate *yr^−^*^1^) (lightning), claycontent, and number of clear days per year (clearDays) were found to be the primary environmental factors associated with decreases in spatial model residual variance (IncNodePurity) and contributed significantly to separating the observations into more homogeneous groups along the trees in the RF model *(SI Appendix, Fig 3a)*. High mean and maximum temperatures also negatively affected the density of tall trees. Tree density increased with photosynthetically active radiation (fapar) by more than 70% but decreased with the number of clear days linked to direct radiation *(SI Appendix, Fig. 4)*. Mean annual precipitation (pannual) and precipitation regimes (pwettest, days20), as well as soil clay content (Claycontet), elevation above sea level (m) (elev), and temperature seasonality (tseason), indicate increases and stability in spatial model-estimated density values *(Appendix, Fig. 5)*. Although water availability is a fundamental resource for the density of tall trees, we observed that poorer or negative relationships for precipitation seasonality (pseason) and potential evapotranspiration (pet) do not reflect an essential dependence across the biome. These results suggest that this set of ecological forces may obscure density trends and patterns at local scales. Despite this counterintuitive observation, the effect of soil water availability (water content) was generally positive or unimodal, decreasing large tree density to a stressed level and increasing after that *(SI Appendix, Fig. 5)*.

### 2.2 Underlying environmental factors

Principal component analysis (PCA) summarizes the complexity of these environmental and ecological relationships across biogeographic space *(SI Appendix, Fig. 6)* and biome (Fig. 3). This approach allowed us to identify the main gradients that drive and shape tall tree density patterns and how these sets of environmental factors correlate. The first two components explained 48.7% of the total variance in the data, with 32.1% attributed to the first axis (Dim1) and 16.6% to the second axis (Dim2) (Fig. 3). In Dim1, the main contributors were precipitation seasonality (pseason, 9.2% contribution) and potential evapotranspiration (pet, 8.7%). These factors point to a solid climatic gradient that separates regions with more significant seasonal variability from areas with more stable conditions. Dim2 highlighted variables related to wind dynamics, such as zonal wind speed (uspeed, 10.1%) and meridional wind speed (vspeed, 9.8%), as well as soil variables, such as clay content (clayContent, 7.5%) and soil water content (waterContent, 7.0%). Lightning frequency also stood out as an essential contributor in Dim2, with a contribution of 6.8%, suggesting its role in local disturbances and mortality processes.

**Figure 3:**
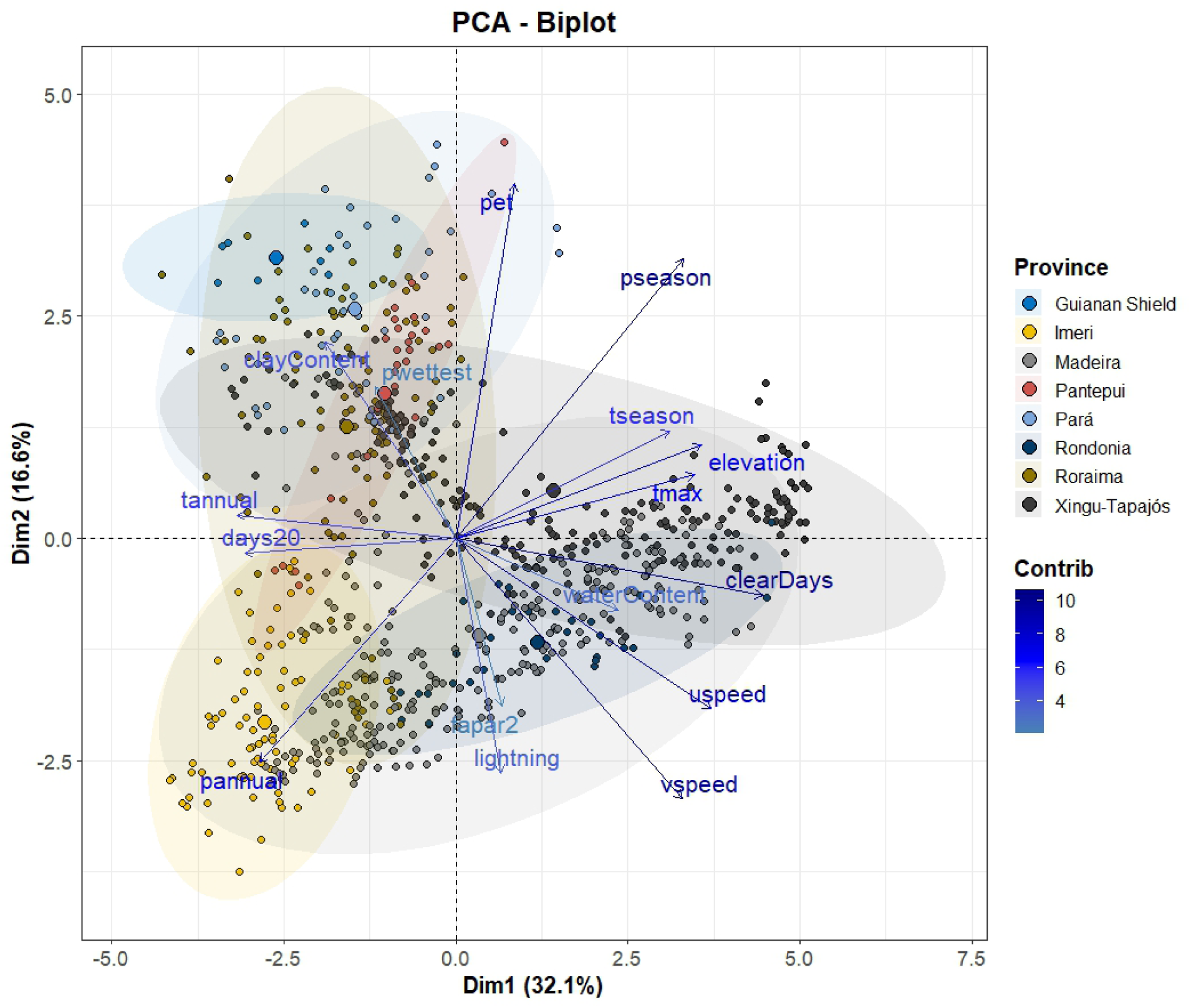
The biplot shows the first two principal components of the PCA, together explaining 48.7% of the data variance. The length of the arrows indicates the degree of contribution (influence) of the environmental variables to the principal axes. Variables with longer arrows (such as pseason, pet, and lightning) significantly impact discriminating biogeographic provinces. The direction of the arrows indicates correlation: Vectors that are close or aligned indicate positive correlation. Vectors that are opposite (*<*90*^◦^*) indicate a negative correlation. For example, pseason positively correlates with Dim1, while uspeed and vspeed correlate negatively. The arrows’ colors reflect each variable’s relative contribution to the model, with a gradient from light blue (low contribution) to dark blue (high contribution). Variables with high contribution best explain the observed variation between provinces, such as pet and lightning. Each point represents a sampling unit (site) within a specific biogeographic province. Ellipses grouped provinces based on their environmental similarities. Example: Guiana Shield (blue) is associated with specific environmental features such as elevation. Pantepui (red) is strongly associated with clayContent and waterContent. The size of the dots reflects the density of tall trees (contribution of emergent species) in each biogeographic province. Areas with larger dots indicate a higher density of tall trees, possibly associated with variables such as lightning and clearDays.

These variables explain the patterns observed in the density of tall trees, evidenced by the size of the points in the graph (Fig. 3). Regions with lower rates of emergent tree density, such as Rondônia and Madeira, presented distinct environmental conditions associated with stronger winds and a more significant number of clear days (clearDays, 6.9%). The Pará and Pantepui provinces showed a lower density of tall trees, probably influenced by poorer soils, extreme climatic seasonality, and stronger zonal winds (vspeed) (see map *SI Appendix, Fig. 4*, and *SI Appendix, Fig. 6)*. In general, the biogeographic provinces were well separated in the PCA space, reflecting differences in environmental conditions that influence the density of emergent trees. For example, Imeri had high rainfall seasonality and higher pet but a low density of emergent trees *(Fig. 3, SI Appendix, Fig. 6)*. These factors indicate that extreme climatic constraints may restrict the growth of emergent trees in this region. With a higher density of tall trees, the Guiana Shield and Roraima were associated soils with higher clay content, less intense winds (uspeed and vspeed), lower frequency of lightning, and clear days. Such conditions may favor seed dispersal and photosynthesis in emergent trees, promoting their growth, establishment, and aggregate density standards. Pará and Pantepui were influenced by edaphic variables such as clayContent and waterContent, suggesting that clay-rich soils with greater water retention capacity support emergent trees in areas with more significant climatic stress. The Xingu-Tapajós province was less correlated with climatic drivers, highlighting intermediate environmental conditions that promote a moderate density of tall trees.

### 2.3 Implications for biomass estimates

At a regional and biogeographic scale, we compared our tall tree density estimates with the potential aboveground biomass (AGB) stock map produced by the same airborne LiDAR campaign as this study (37). As expected, higher density of tall trees is strongly associated with higher aboveground biomass (R = 0.5, p *<* 2.2e16) (Fig. 4a). These relationships are more pronounced where large tree density is higher, specifically in the provinces of Roraima and Guiana Shield, which also correspond to the largest AGB densities across the biome. Interestingly, Xingu-Tapajós and Madeira have relatively high AGB even in areas with low density of tall trees. However, in areas with higher densities of tall trees the relationship is not linear and may indicate saturation at extreme density rates. For example, areas with low density of tall trees may still have significant biomass levels due to the contribution of smaller trees. The relationship between giant tree density and aboveground biomass was also analyzed in different classes of tree density (0–50, 50–100, and 100–150 trees/km2) (Fig. 4b). We observed that, across the biome, spatial patterns agree well and that our density estimates are predictive of biomass even in biogeographic provinces with densities of up to 50 tall trees/km2. Notably, the relationship between biomass and density is most significant in areas with more than 100 tall trees/km2 (Fig. 4b). The results reveal a consistent increase in biomass with increasing tall tree density, mainly in Roraima and the Guiana Shield. The classes with the highest density (100–150 trees/km2) presented the highest median biomass values (40.0–50.0 Gt/km2) and the lowest variability, while the lower density classes (0–50 and 50–100 trees/km2) exhibited more visible dispersion in biomass values, reflecting structural heterogeneity.

**Figure 4:**
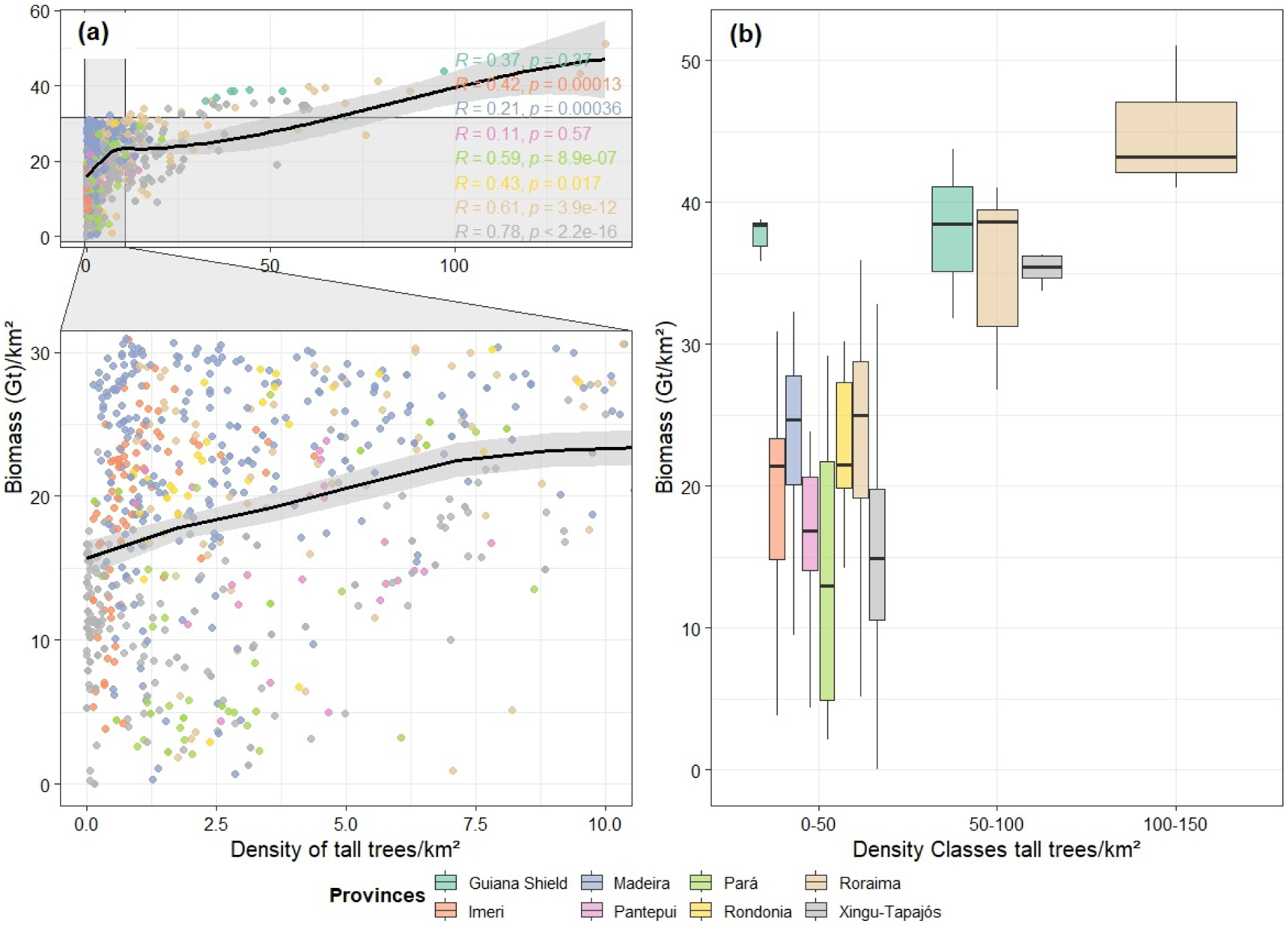
Estimated aboveground biomass in relation to the density of tall trees (i.e., ⩾ 60 m) for the 900 LiDAR plots studied in the Amazon biome. (a) Scatterplot with all the 900 points, (b) Cropped area showing the relationship for areas with densities *<* 10 tall trees km2; (c) biomass per tall-tree density class. The graph’s black line represents the relationship’s goodness of fit (linear model - method = ‘less’). At the same time, the gray bands around it indicate the confidence interval, suggesting a significantly feasible overall relationship (R = 0.5, p *<* 2.2e16). The graph highlights the significant contribution of emergent trees to biomass and carbon stocks in the Amazon.

## 3 Discussion

### 3.1 Patterns of tall tree density and environmental drivers

The density of tall trees is a valuable input for modeling large-scale biological and biogeochemical processes and a prominent component of ecosystem structure, governing elementary processes such as retention rates, competitive dynamics, and habitat suitability for many plant and animal species (1, 4, 9). Our results show that a complex combination of environmental factors influences the density of tall trees in the Amazon. Clay-rich soils, moderate precipitation and temperature, and low incidence of lightning and strong winds create stable conditions for a high density of tall trees. These findings are consistent with the existing literature highlighting the importance of water availability and stable climatic conditions for the growth and maintenance of large trees (28, 39). Areas located in the northeastern portion of the biome, between the provinces of Roraima and the Guiana Shield (approximately -58*^◦^* to -51*^◦^* longitude, and -4*^◦^* to -3*^◦^* latitude), have the highest local density of tall trees across the Amazon. We estimate up to 140 individuals per km2 in these areas, where well-drained hilltops terra firme forests predominate. These patterns confirmed previous results at the same scale for biomass (37), maximum height (28), and diversity of large species (31). Furthermore, these areas are among the most remote and least explored in the Amazon (33), suggesting a possible role of a low current human population density and the absence of roads or large infrastructures as factors controlling the occurrence of tall trees. Being isolated areas with low anthropic pressure enhances the conservation value of these areas.

Lightning and strong winds associated with convective storms and cold fronts in the southern and western Amazon can cause significant mortality of tall trees through severe damage, toppling, or falling (40–42). Tall trees that emerge above the forest canopy are particularly susceptible to these disturbances, as their height exposes them to greater lightning strikes and the risk of falling caused by strong winds (41–43). Recurrent increases in wind speed can significantly modify density patterns in many regions (44). Recent research shows that lightning has been a prominent cause of canopy damage in the tropics, often leading to tree mortality and consequent reduction in large tree density (22, 45, 46). Since most trees in the biome are not tall (average canopy height 30–45 m, e.g., Lang et al.(47)), lightning and wind are much less critical for forests with lower canopies (48).

At the local scale, other factors become more relevant in explaining the variation in the density of tall trees. Topography, treated here as a proxy for altitude, was fundamental. The tall trees are often associated with plateaus or higher areas (16, 49). In contrast, the aggregation of smaller trees tends to occur in lowlands or terrains with more significant soil heterogeneity (31). Studies, such as that of Zuleta et al. (49), highlight that uneven terrain increases the spatial aggregation of trees of similar diameters, contributing to local density patterns. Low-lying areas (*<* 50 m above sea level) in the Amazon, especially in ecotone regions such as the Imeri province, are associated with lower densities of tall trees due to a combination of ecological and ecophysiological factors. Low altitude generally coincides with flat terrain and proximity to river systems, where permanent, seasonal, or transient flooding occurs (50–52). This generates unfavorable hydrological conditions for the growth of large trees, affecting both the availability of oxygen in the roots and the nutritional stability of the soil (53, 54), which are fundamental for the development of tall trees. In these conditions, species prioritize adaptations that favor survival and tolerance to flooding rather than investing in vertical growth (55, 56). This can explains why forest communities in low-lying and flooded regions generally comprise taxa that do not reach great heights and canopies (57). The vegetation in these ecosystems is often exposed to flooding regimes that impose intermittent water stresses (52, 58), and favour species with smaller height and crowns that can maintain higher mechanical and physiological stability (54, 59, 60). In addition, the variation in sediment and nutrient loads from rivers in these areas can contribute to a higher instability in nutrient availability, which compromises the structural development necessary for trees to reach great heights (54, 60). Therefore, recurring and seasonal water stress on the root system can compromise the roots’ ability to capture oxygen, impairing cellular respiration and tree growth (61, 62). This limitation in respiration is particularly detrimental to tall trees, which require a significant amount of energy to sustain processes of transporting water and nutrients over large vertical distances (63–65). In conditions of low oxygenation, roots cannot sustain the efficiency necessary to supply resources to upper tissues, limiting growth potential (54, 66).

Our findings also indicate that clay and soil water content are critical for the occurrence and density of giant trees in the Amazon, as they directly influence nutrient availability and water stability, which are essential for the growth of these trees (67, 68). Soils with high clay content have a higher cation exchange capacity, which allows the retention of essential nutrients such as calcium, magnesium, and potassium, favoring the sustained growth of trees. In addition, clay contributes to the formation of aggregates in the soil, improving its structure and increasing water retention. In Amazonian regions with clayey soils, giant trees’ density and biomass are higher than in areas with sandy soils due to their water retention capacity (56). Soil water availability, especially during dry periods, is crucial for the survival of these trees, and clayey soils provide a stable source of moisture. However, other studies suggest that in the tropics, higher soil fertility is linked to lower canopy height (69, 70). Plants increase their productivity and tend to die younger because they have invested all their energy in growth rather than defense and longevity. For example, plants of the same species in fertile soils tend to grow faster and have lower wood density, so they are more likely to break in stronger winds (23, 56, 71, 72).

In environments well drained with regular rainfall and defined seasons, there is a constant supply of water in the soil, which is essential for continued growth and for the maintenance of complex physiological processes that require high levels of water, such as transpiration and photosynthesis for many tall trees *(SI Appendix, Fig. 3b)*, and this is directly related to the number of days with clouds or days with few clouds (”clear days”). An increase in cloudless days indicates a greater incidence of sunlight and high temperatures, which can increase the rate of photosynthesis, favoring tree growth to a thermal tolerance threshold (73, 74). Although atmospheric drought, caused by the combination of these factors, can increase photosynthetic activity in the Amazon rainforest when water supply is abundant (73, 75), we can deduce that in long periods of drought, tall trees may suffer more easily from embolism (76–78), increasing hydraulic vulnerability to water stress due to prolonged exposure of plants to high temperatures with intense solar radiation and associated prolonged periods of seasonal water deficit (24). Bennet et al.(13) found that droughts consistently negatively impacted the growth of larger trees and mortality rates on a global scale. Other studies may suggest that the number of clear days and seasonality of precipitation are drivers of many density patterns on a regional and continental scale (79–82).

### 3.2 Importance of tall trees density for biomass stocks

The positive relationship between tall tree density and biomass for the entire biome and biogeographic provinces highlights the importance of emergent trees, which play an essential role in biogeochemical processes despite representing a small fraction of the total trees (2, 14). For example, in the tropics, Sullivan et al. (74) showed that only 1% of trees in tropical forests account for about 50% of aboveground biomass. Slik et al. (4, 83) suggest that the impact of large trees on AGB is, on average, 25.1% in South America and more than two-thirds at the pantropical scale. These results are consistent with the meta-analysis by Lima et al. (84) in the Amazon biome, which highlighted that only one giant tree could accumulate 82% of all stored AGB *Mg ha^−^*^1^. In general, tall tree density maps can potentially characterize habitat heterogeneity directly (1, 85), so canopy height has been classified as a high-priority biodiversity variable to be observed from space (47). Therefore, the density of tall trees can be used as a direct indicator of the potential for carbon storage in tropical forests.

Our study did not test the effect of local processes influencing the density of tall trees on aboveground biomass. However, a possible explanation for the high AGB in biogeographic provinces dominated by large trees may be related to the reproductive strategy of wind dispersal of some species, i.e., they need to be tall and emergent to effectively disperse their seeds and exceptionally maximize their geographic influence (4, 86). In the Amazon, the most abundant larger species (dbh ⩾ 70 cm) were *Goupia glabra*, *Dinizia excelsa*, *Aspidosperma excelsum*, *Couratari guianensis*, *Manilkara huberi*, *Dipteryx odorata*, *Tabebuia serratifolia*, *Bertholletia excelsa*, and *Caryocar villosum* (84). These species represented approximately 20% of the individuals sampled in 240 plots inventoried in the eight biogeographic provinces in the biome. Therefore, although environmental controls influence maximum tree height and the diversity of large species at a regional scale (28, 31), this dominance is also driven by species-specific traits that provide the ability to reproduce and support large biomass stocks at a local scale (87). Different strategies in seed dispersal are linked to maximum tree height and aggregation patterns (86, 88, 89). In particular, wind-dispersed species such as *Dinizia excelsa*, *Aspidosperma excelsum*, and *Tabebuia serratifolia* show more aggregated patterns than animal-dispersed species such as *Bertholletia excelsa* and *Manilkara huberi*, which may affect large tree density and AGB at the local scale, as animals are more effective at dispersing seeds in a more uniform spatial distribution (90). This finding emphasizes the importance of including species traits and changes in species composition as explanations for AGB gradients in tropical forests (4, 83, 87).

Another critical factor at the local scale is gap formation, which provides opportunities for seed germination and recruitment, especially in areas where light is a limiting resource. Higher densities of tall trees produce more significant gaps when they fall (91, 92), although the frequency and size of these gaps are also influenced by disturbances such as wind and lightning (41, 93, 94). A relatively low proportion of significant gaps is observed in the provinces of Roraima and Guiana Shield, where we found the highest densities of tall trees and stored biomass (probably caused by the low mortality rate of large trees caused by strong winds and lightning). These results suggest that low gaps may limit the recruitment of tall trees, contributing to their aggregated patterns and more prominent AGB stock over time, as germination and recruitment may be limited to fewer areas with more significant gaps. For example, clusters of *Dinizia excelsa* and *Tabebuia serratifolia* can be found in an aggregated form, likely due to a combination of anemochoric seed dispersal, a potential dependence on light for germination and seed recruitment, as well as local topography (lowlands versus plateaus) that favor their aggregation patterns (84, 95, 96).

The results presented here reinforce the hypothesis that preserving areas with a high density of emergent trees is crucial to maintaining ecosystem services associated with carbon storage. Including tall tree density as a critical variable in biomass estimation models can substantially improve the accuracy of these estimates. These models need to better account for the role of these large trees and airborne LiDAR surveys is a practical tool detect and quantify large trees and quantify their contributions to biomass stocks (34, 36). This is especially interesting in difficult-to-access regions with limited ground-based measurements. The observed relationship suggests that the loss of tall trees due to deforestation or forest degradation can significantly reduce biomass stocks. Inadequate management that does not consider the conservation of these trees can have drastic implications for the carbon budget, amplifying the impacts of greenhouse gas emissions.

### 3.3 Limitations of the study

Firstly, linking density data with variables such as wind, lightning, solar radiation, precipitation, and soil clay and water contents can present challenges due to the complexity of ecological interactions. Models that attempt to predict density based on these variables may only capture some of the nuances of these interactions, leading to potentially inaccurate inferences (97).

Secondly, *SI Appendix, Fig. 2(a)* illustrates the distribution of the tall tree density across the mapped transects. Most transects exhibit low-density values, while only a few exhibit high-density. This unbalanced sample distribution has the potential to introduce bias into model training, given the tendency of the Random Forest algorithm to be more influenced by frequent data patterns. *SI Appendix, Fig. 2(b)* presents the relationship between observed and predicted values by the Random Forest model, demonstrating good overall predictive ability. However, it is essential to note that there are discrepancies at the extremes. The model predictions for low-density transects are more numerous, potentially leading to a lower root mean square error (RMSE) in these areas. On the other hand, the RMSE for high-density plots is likely higher due to the smaller number of samples available to represent these cases in the model. This significant sampling bias can impact the spatial model results, as areas with low density of tall trees are better represented and therefore better predicted than areas with high density. This inequality can result in a model that either underestimates or overestimates density in less-represented areas. To mitigate this effect, we consider data balancing techniques or methodological adjustments that directly address variability at different density levels through spatial cross-validation, thereby improving model robustness and accuracy. Although attempts have been made to correct density values for these sampling biases, they likely influenced, at least partially, the results.

As a final point, it is crucial to highlight that different forms of land use significantly contribute to high deforestation rates, although these are not considered variables in this study. This drives reductions in the density of tall trees, especially in the southern Amazon, and can obscure the expected effects of climate, topography, and soil in several regions with tall trees. As a result, forest ecosystems are often relegated to drier regions, reversing expected within-biome relationships between moisture availability and tree density (11). Such effects may vary across biogeographic provinces, states, and municipalities depending on human population density, alternative resource availability, and socioeconomic status. Therefore, future studies should consider these factors to provide a more comprehensive understanding of the dynamics of tree density in the Amazon biome.

### 3.4 Broader implications and concluding observations

The main findings of this study are significant for understanding the ecology and conservation of giant trees in the Amazon biome. Mapping tall tree density provides valuable insights into forest structure and health, which are crucial for biodiversity conservation and climate change mitigation. The data generated can inform forest management and conservation strategies, helping to identify areas in need of protection or restoration. Furthermore, by linking tree density to environmental variables, the study can reveal the key factors influencing the distribution and health of tall trees, and offers a beacon of hope in the face of climate change and deforestation, providing a roadmap for more effective conservation efforts. Practical application of the data must be accompanied by a deep understanding of ecological interactions and an adaptive approach to environmental management.

The relationship between tall tree density and local factors, such as soil characteristics, dispersal strategies, and gap formation is a promising area for future investigation. The hypothesis that local density correlates with reproductive success and recruitment can be tested in different soil types and terrains. Furthermore, exploring density patterns across topographic gradients and relating them to climatic and edaphic variables can provide insights into the interaction between regional and local factors. Regional factors, such as wind and lightning, shape tall tree density patterns in the Amazon. However, they are refined at local scales by processes such as dispersal, topography, and gap formation. These results highlight the importance of integrating multiple scales of analysis to understand the ecology of giant trees in the Amazon and provide support for their conservation under a climate change scenario.

This study contributes to the discussion on the relevance of approaches based on high-resolution data, such as those provided by LiDAR sensors, to better understand tropical forests’ structural dynamics. Furthermore, combining tall tree density data with climate change projections could provide insights into the future of carbon storage in the Amazon. Considering their disproportionate importance in biomass stocks, conserving tall trees should be a priority in global conservation initiatives such as REDD+ (Reducing Emissions from Deforestation and Forest Degradation). Investments in remote monitoring and forest inventories that integrate metrics from emerging trees can improve understanding of forest dynamics and support global mitigation strategies.

Although this study does not aim to model the potential impacts of climate change, it is concerning to note that several climate variables strongly associated with giant tree density may undergo significant changes in future climate crisis scenarios. Environmental changes induced by rising temperatures are already being observed. Changes in environmental variables associated with disturbances can significantly negatively impact the density and survival of large trees. For example, the frequency of anomalous events, such as increased storms and lightning strikes, has already been observed (98–101). Furthermore, in the current scenarios projected by the sixth climate report of the Intergovernmental Panel on Climate Change (IPCC), the average global temperature is estimated to increase by 1.5°C between 2030 and 2052 and, if it continues to increase and precipitation levels decrease, and drought scenarios in the Amazon will sadly become more frequent, directly impacting all biodiversity (102). Even in this scenario, only approximately 15% of the Brazilian Amazon is protected by conservation units, covering only about 58% of the remaining vegetation. Given the ecological importance of giant trees, understanding the effects of climate change on density patterns is critical and it should be explored quickly at finer scales within the Amazon. This knowledge is critical to refining conservation perspectives in a changing world—for example, to what extent are protected areas in the Amazon currently susceptible to impacts induced by climate disruptions, and how can the giant trees withstand or respond to these changes? Efforts to understand how deforestation and climate change interact and mitigate their impacts are urgently needed in light of the high and increasing rates of deforestation in the Brazilian Amazon in recent, and that directly threaten sanctuaries of ancient trees throughout the biome.

## 4 Materials and Methods

### 4.1 LiDAR data collection and standardization

The point cloud from airborne LiDAR was acquired between 2016 and 2018 for 900 transects distributed throughout the Brazilian Amazon biome (Fig. 1). The data collection was part of the EBA (Biomass Estimation in the Amazon) project developed by the National Institute for Space Research (28, 37). The sensor used was the LIDAR HARRIER 68i, coupled to a CESSNA model 206 aircraft. The scanning angle was 45 degrees and the flight altitude was approximately 600 meters. The point cloud is formed by 4 returns per square meter. Each transect mapped 375 hectares (representing 12.5 x 0.3 km strips). See more details at zenodo.org.

For each transect, an approach based on local maxima (canopy height model – CHM) was applied to identify tall trees using a moving window of 100 meters (103). The located treetops were grouped into three vertical strata: heights *<* 40 m; with heights ⩾ 40 m and *<* 60 m; and heights ⩾ 60 m. A unique string for each transect assigned as ID (transect number); the transect central coordinates (LAT and LON) and total area (in hectares) formed the initial dataframe for subsequent analyses (Supplementary Table 1). To ensure consistency and maximum accuracy in the density data per transect and the final density maps, we standardized the area size to square kilometers (km2). Considering only the tall trees (height ⩾ 60 m), we found 313 transects (≈ 35%) distributed throughout the Amazon biome to further analysis.

All transects in the final tree density data matrix cover eight biogeographic provinces proposed by Morrone (104). This biogeographic definition seeks a universal classification in provinces with similar macroecological characteristics of biodiversity. To understand the optimal environmental conditions for occurrence of the tallest trees, the density data per km2 were linked to spatially explicit environmental factors. The wide distribution of our sampling points (in number and distribution of LiDAR transects) provides a substantially representative sampling effort for this vegetation stratum. It ensures that any uncertainty in transect locations or minor changes in forest area in the Amazon region are unlikely to alter mean values or estimates of giant’s tree density.

### 4.2 Environmental factors

Our research involved initially 16 spatially explicit environmental predictor variables representing topography, climate, and soil, as detailed in *SI Appendix, Table 1*. The data were then carefully cropped to fit the geographic boundaries of the Brazilian Amazon biome and, when necessary, resampled to a spatial resolution of 30 arc seconds (≈ 1 km).

Information on temperature and precipitation derived from 19 bioclimatic variables was obtained from WorldClim version 2 (105). The average number of cloudless days throughout the year was obtained using surface reflectance products from the MODIS (Moderate Resolution Imaging Spectroradiometer) sensor. We used the Terra MOD09GA Version 6 product, which estimates the MODIS surface spectral reflectance duly corrected for atmospheric conditions.

The annual average number of days with precipitation greater than 20 mm was calculated from the precipitation time series of the Climate Hazards Group InfraRed Precipitation with Station (CHIRPS) dataset (106). Potential evapotranspiration was derived from data provided by TerraClimate, which combines WorldClim climatological normals, Climatic Research Unit (CRU) Ts4.0, 55-year Japanese Reanalysis (JRA-55) data, and the Penman-Monteith methodology. The fraction of absorbed photosynthetically active radiation (FAPAR) was obtained from the calibrated and corrected land surface reflectance product of the Advanced Very High-Resolution Radiometer (AVHRR) of the National Oceanic and Atmospheric Administration (NOAA), providing information on the photosynthetic activity of plants (107).

Lightning frequency, associated with weather events and tree mortality (45), was obtained from the Lightning Imaging Sensor (LIS) instrument aboard the Tropical Rainfall Measurement Mission, provided by NASA’s Earth Observing System’s Global Hydrological Resources Center (EOSDIS). Wind speed data were provided regarding the five-year average daily maximum speeds from the fifth global model reanalysis (ERA5) of the European Centre for Medium-Range Weather Forecasts (ECMWF).

Two wind speed metrics were used: u-speed, representing the zonal (eastward) component, and v-speed, representing the meridional (northward) component. Previous studies indicate that winds are correlated with disturbances that result in tree mortality in the Amazon (43, 44).

Soil variables were obtained from SoilGrids, based on the World Reference Base (WRB) and USDA classification systems, totaling approximately 280 raster layers (108). The layers referring to clay content (% fine particles *<* 2 *µ*m) and water (% volume at field capacity in 30 cm), both with a spatial resolution of 250 m, were estimated through machine learning applied to a compilation of global profiles and soil layers (108). All geospatial data were processed in ArcMap 10.1 software. We extracted all geospatial covariate values from raster datasets for transect location points using the raster::extract function from the R raster package (109) to construct a standardized plot-level dataframe.

### 4.3 Spatial modeling

A random forest model was used to model the density of tall trees from the environmental variables (110). This machine learning method detects global trends present in data using a substantial ensemble of decision trees to predict tree density in the uppermost strata of the Amazon rainforest canopy using the 16 environmental covariates. The RF algorithm applies the general bootstrap aggregation technique (bagging) with a modified tree learning algorithm that selects a random subset of the features at each candidate split in the learning process (111). Since a random subset of variables is chosen for each tree, the RF algorithm based on trained tree ensembles avoids overfitting (112). It circumvents potential multicollinearity issues (113) between the predictor variables.

For rigorous evaluation of the RF model, we employed the k-fold cross-validation method and spatial cross-validation. Firstly, The 313 sampled transects were randomly divided into “k” subsets (or “folds”) of approximately equal size. The randomized cross-validation was of the k-fold type, randomly dividing into “k” groups. In this procedure, “k” is defined as 15. For each “k” subset, the Random Forest model is trained using k1 subsets as the training set and the remaining subset as the test (or validation) set. This process was repeated 100 times with sample replacement to examine the accuracy of the estimated tree density values. After each round of training and validation, we calculated model performance metrics, such as root mean square error (RMSE) and coefficient of determination (R2). These metrics are stored for each fold. At the end of the process, the metrics obtained in each “k” validation round are aggregated, usually by the average, to provide an overall estimate of the model performance. This average provides a more stable and reliable measure of model accuracy than a simple single split between training and testing.

A spatial cross-validation methodology was employed to assess the predictive ability and spatial uncertainty of a Random Forest (RF) model in estimating the density of tall trees (height ⩾ 60 meters) based on environmental variables. The spatial cross-validation approach is particularly suitable for models applied to spatial data, as it accounts for the spatial autocorrelation among sampling points, thereby minimizing the overestimation bias of model accuracy metrics due to the spatial proximity between training and test data. Initially, the data were organized in a spatial structure using the sf library (114), allowing for the manipulation and visualization of sample points with geographic coordinates. Ten folds were used for spatial cross-validation, each representing a unique data partition. This division was conducted randomly and in a balanced manner, ensuring that each sample point was used exactly once as a test set. In contrast, the others were used for training, thus ensuring the generalization of the results and control over prediction variability.

For each fold, the model was fitted using the ranger package (115), an efficient implementation of Random Forest configured with 500 trees. The response variable was the density of tall trees (N trees h60 km2). The explanatory variables included climatic, topographic, and edaphic environmental factors, carefully selected to capture spatial variability in environmental conditions. To avoid missing data issues (NA), the complete cases function was applied to the training data, ensuring that only complete records were used to fit the model. After fitting the model in each fold, predictions were made for the corresponding test set. These predictions were stored in a matrix, where each column represented the predictions of a specific fold. The uncertainty associated with the predictions was estimated by calculating the standard deviation across folds for each sample point, creating an uncertainty map. This standard deviation measure reflects the variability in predictions across different folds, indicating areas with greater or lesser confidence in the model estimates.

The statistical metrics used to evaluate model performance included the root mean square error (RMSE) and the coefficient of determination (R2) of the tall tree density calculated for each fold. RMSE is a measure of accuracy that reflects the average magnitude of prediction errors, while R2 represents the proportion of variance explained by the model. Both provide a comprehensive assessment of model accuracy. Additionally, the mean bias was calculated to identify potential systematic deviations in predictions, offering an additional metric to verify the spatial accuracy of the model.

After cross-validation, the final model is fitted using the entire available dataset (all 313 transects) and is used to predict giant tree density across the study area, considering environmental factors as predictor variables. The importance of environmental variables was analyzed using marginal plots, holding the other variables constant at a mean value. This approach consisted of a sensitivity analysis in which the importance of variables is measured by permuting variables in the model and measuring the increase or decrease in tree density. Finally, the parameters of the final RF model were applied to stacked environmental layers at the pixel level for the entire Amazon biome using map algebra to produce density maps for the uppermost strata of the Amazon forest canopy, especially giant trees *(SI Appendix, Fig. 1)*. All statistical and spatial modeling and analysis procedures were conducted in the R environment version 4.2.1 (116), using the MASS (117) and randomForest (110) packages.

### 4.4 Principal component analyses

To investigate the correlation patterns between giant tree density and environmental factors, we implemented principal component analysis (PCA). PCA was conducted separately for each biogeographic province and the Amazon biome. Biplots were generated for each case showing the first principal components using the factoextra R package (118), searching for local and regional patterns. This statistical technique allowed us to identify the main linear combinations of environmental variables that explain most of the variance in the data and also to understand how environmental factors vary between and within each province, indicating that the factors driving giant tree density patterns may be regional and highly influenced by local environmental conditions. Prior to analysis, the data were normalized to avoid biases associated with different scales between variables, applying the following steps:

The data were normalized to avoid biases arising from different scales. Normalization was performed using the standard transformation:

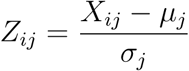

where *Z_ij_*is the normalized value of the *i*-th observation of the *j*-th variable, *X_ij_*is the original value, *µ_j_* is the mean of variable *j*, and *σ_j_* is the standard deviation of *j*.

PCA was applied based on the covariance matrix of the normalized data. The covariance matrix was calculated as:

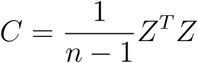

where *C* is the covariance matrix, *Z* is the normalized data matrix, and *n* is the number of observations.

The principal components (PC*_k_*) were obtained by solving the eigenvalue problem:

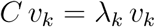

where *v_k_*are the eigenvectors (directions of the principal components) and *λ_k_* are the eigenvalues (variance explained by each component).

The principal component analysis allowed us to extract orthogonal axes of environmental variation, providing a quantitative basis for exploring how environmental factors correlate within each province and how these correlations drive giant tree density patterns in the Amazon biome.

### 4.5 Assessing biomass estimates

To estimate above-ground biomass (AGB) as a function of tall tree density in the Amazon biome, we used a reference dataset from a raster biomass map developed by Ometto et al. (37). This map was generated from the same 900 flyovers randomly distributed throughout the Amazon biome and its biogeographic provinces, ensuring compatibility and spatial representativeness of the data. The original biomass map, with a spatial resolution of 50 meters, was resampled using the nearest neighbor method to adjust its resolution to 250 meters. This procedure was necessary to ensure compatibility with the map of tall tree density, also derived from the same 900 flyovers. The extraction of biomass values per transect was performed using the raster::extract function of the raster package. After extraction, the values were converted from Mg/ha to Gt/km2 to allow a standardized interpretation of biomass stocks at a spatial scale.

With the processed data, we evaluated how the density of tall trees can explain biomass stocks in the Amazon biome and its biogeographic provinces. To do so, we fitted a linear model of the type *AGB = density of tall trees*. This model allowed us to identify the relationship between the density of emergent trees and biomass stocks, considering variations at both the biome and biogeographic province scales. This methodological approach enables a robust analysis of the role of tall trees in biomass stocks in the Amazon, contributing to the understanding of the spatial distribution of carbon stored in tropical forests and the factors that influence this dynamic at different geographic scales.

## Authors’ contributions

R.B.L designed research; R.B.L, D.A.S, M.H.N, P.R.L.B, P.G, C.P.O, T.J, D.G.S and E.B.G. performed research; R.B.L, M.H.N, T.J, N.H, D.R.A.A, J.P.O and E.B.G. provided funding; R.B.L, CPO, and EBG. analyzed data; and R.B.L, M.H.N, P.R.L.B, CPO, D.A.S, PG, and EBG wrote the paper and all authors edited the manuscript.

## Acknowledgements

Funding was provided by the Coordenação de Aperfeiçoamento de Pessoal de Nível Superior Brasil (CAPES; Finance Code 001); Conselho Nacional de Desenvolvimento Científico e Tecnológico (Processes 403297/2016-8, 301432/2022-8, 301661/2019-7, and Processes 444350/2024-1); Amazon Fund (grant 14.2.0929.1); National Academy of Sciences and US Agency for International Development (grant AID-OAA-A-11–00012); Universidade Federal dos Vales do Jequitinhonha e Mucuri (UFVJM); and Instituto Nacional de Pesquisas Espaciais (INPE). R.de Lima was supported by the Universidade do Estado do Amapá (Processes 0022.0279.1202.0018/2021). P. Groenendijk was financial support by the São Paulo Research Foudation - FAPESP, Young Investigator Grant (2018/01847-0).D. Almeida was supported by the São Paulo Research Foundation (2018/21338-3; 2019/14697-0). N. Higuchi and P. Aparicio were suppor ted by INCT-Madeiras da Amazônia and Next Generation Ecosystem Experiments-Tropics (NGEE-Tropics), as part of DOE’s Terrestrial Ecosystem Science Program Contract No. DE-AC02-05CH11231. T. Jackson was supported by the UK Natural Environment Research Council grant NE/ S010750/1. M. Nunes was supported by the NASA – University of Maryland “Global Ecosystems Dynamics Investigation” (GEDI) spaceborne LiDAR mission.

## 5 Supporting Information

**Figure S1:**
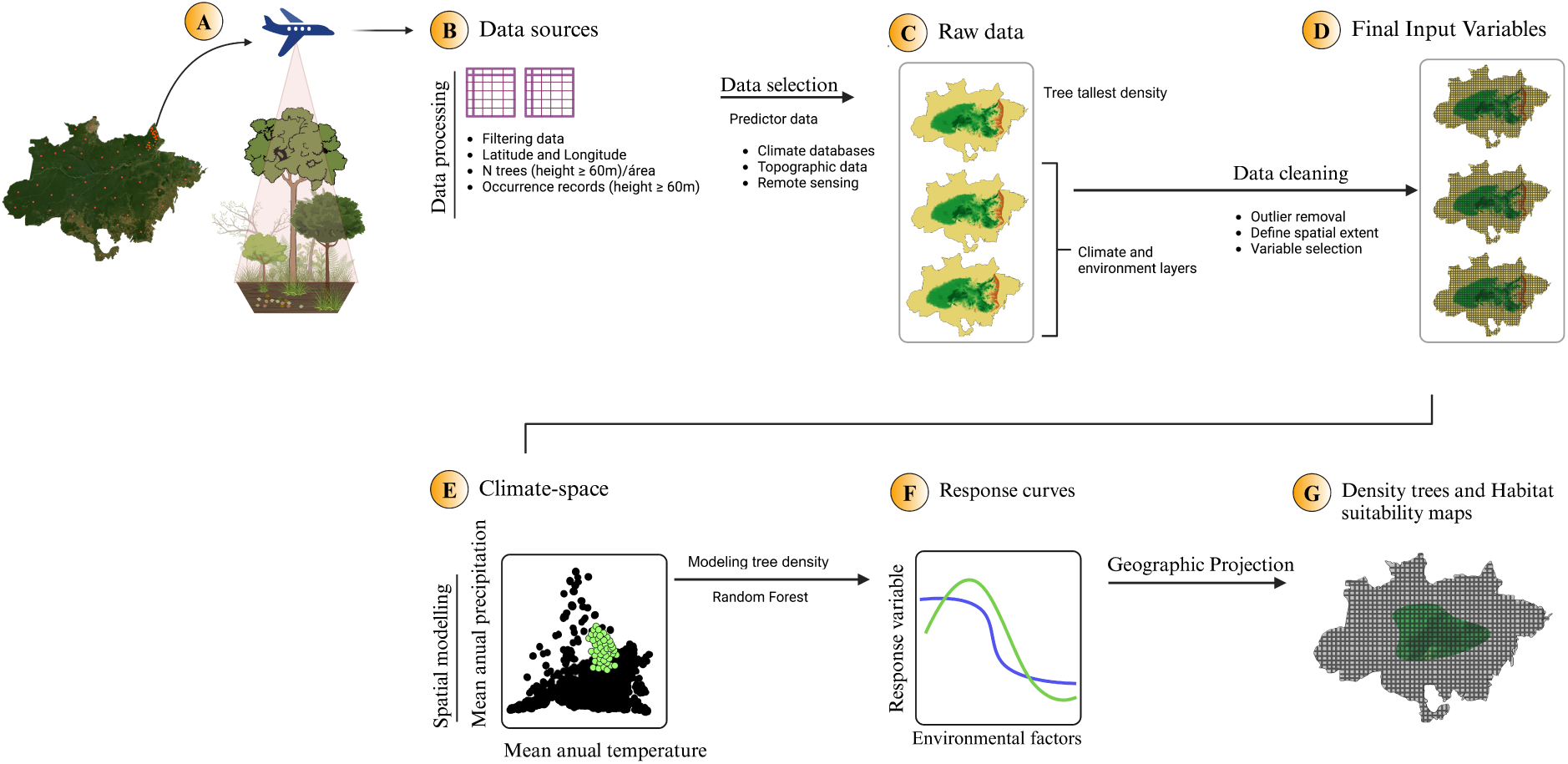
Methodological flowchart for mapping giant trees in the Amazon, highlighting data collection, processing, and modeling steps. (A) Data collection: airborne LiDAR surveys are employed to identify and record giant trees (height ⩾ 60 m) and location data, ensuring the accuracy of our findings in the Amazon. (B) Data sources and processing: Occurrence data (trees ⩾ 60 m) are meticulously filtered based on latitude, longitude, and tree density per area. (C) Raw data: Comprehensive climate, topographic, and remote sensing databases are used to generate environmental layers and density records of giant trees (D) Final input variables: Data cleaning involves removing outliers, defining the spatial extent, and variable selection for modeling purposes. (E) Spatial tree density modeling by RF model using a climate variable space (e.g., mean annual precipitation and temperature) and other factors. (F) Response curves: Response curves relate the density of giant trees to environmental variables, showing how these variables influence the density. (G) Density maps: Geographic projection of the modeling results, producing maps that show the density of giant trees across the Amazon. The figure presents an analysis of the density of tallest trees in the Amazon basin, a study that unveils crucial insights into the distribution of observed and estimated data and the effectiveness of the spatial model generated by the Random Forest algorithm. These results are of paramount importance in understanding the predicting spatial model of the tallest trees at Amazon basin’s.

**Figure S2:**
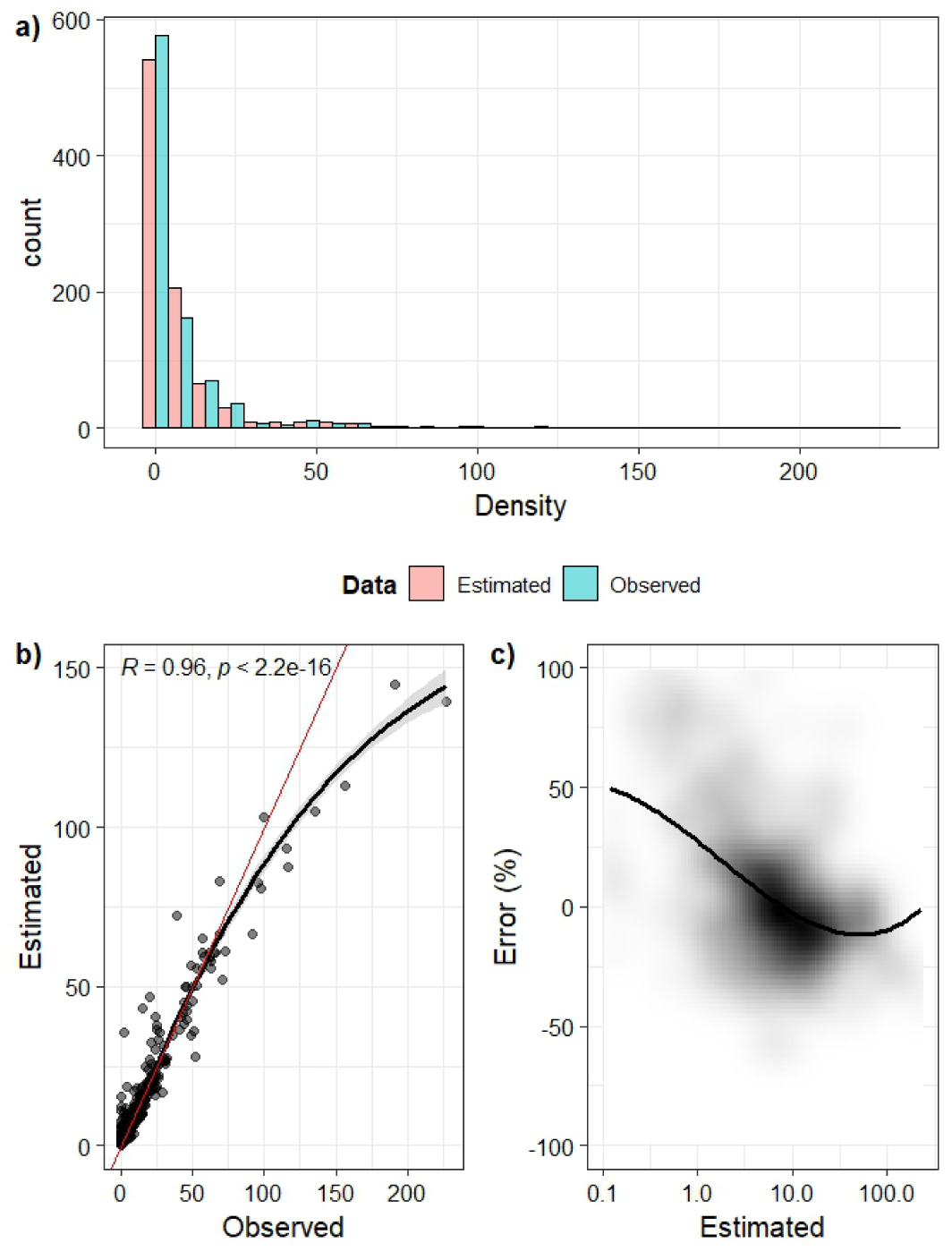
Distribution and analysis of the density of tall trees in the Amazon basin. Graph a) shows the observed versus estimated data distribution, highlighting a distortion in the curve’s tail that suggests underestimation in places with high tree density. Graph b) presents the prediction of density values by the Random Forest spatial model. Graph c) indicates the residual distribution in percentage, highlighting the effectiveness and limitations of the model in spatial prediction. Comparing the distributions of observed data with the values estimated by the model, we note an evident distortion in the curve’s tail, especially in regions with a high density of tallest trees. This distortion suggests that the model may be underestimating the number of trees in areas with a high concentration of the tallest trees. This underestimation can be considered critical, as it may indicate that the spatial model is not adequately capturing the variables that influence maximum tree density. Graph b) illustrates the prediction of the highest tree density values generated by the Random Forest spatial model. Despite being robust and widely used in spatial analysis, the model presents more significant errors in accurately predicting areas of high density, evident by the distortion observed in graph a). This suggests that, although the model may be effective in predicting medium or low tree density, it requires adjustments to deal with the extremes of the distribution. Graph c) presents the distribution of residuals in percentage, which was calculated by subtracting the observed values from the values predicted by the model. The analysis of the residuals reveals that, although the model is relatively accurate in most areas studied, there is a significant concentration of errors in areas with a high density of taller trees. When analyzed, these positive and negative residuals provide critical insight into the model’s limitations in capturing spatial variability in extreme regions of the Amazon basin. The analysis of the essential variables for estimating the density of the tallest trees in the Amazon basin, represented in the figure, provides a clear view of which environmental factors are most influential in the spatial model generated by Random Forest.

**Figure S3:**
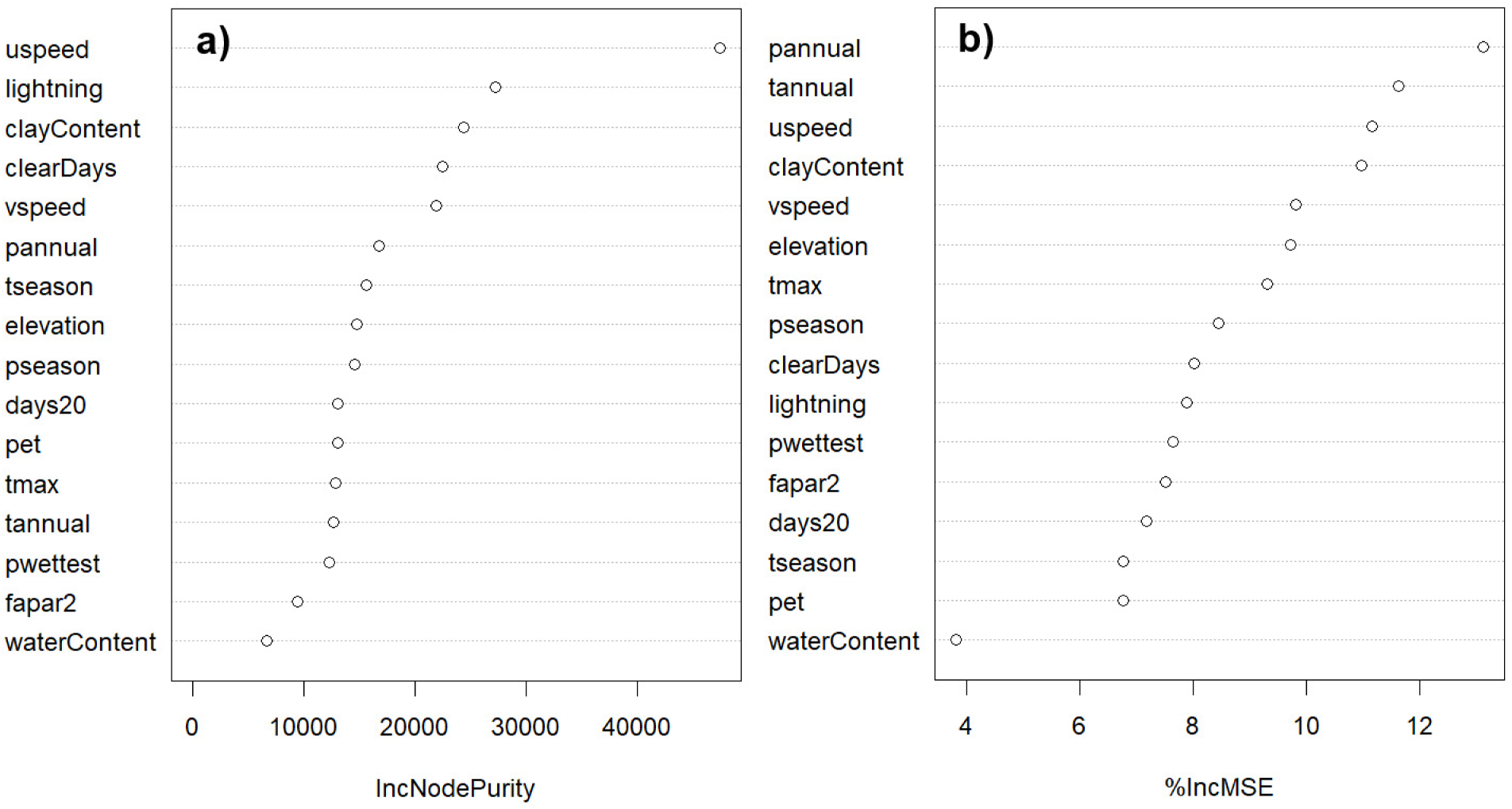
The Crucial Variables in Estimating the Density of the Tallest Trees in the Amazon Basin. Graph a) showcases the most critical variables, as measured by the IncNodePurity metric, underscoring the substantial influence of topographic and climatic variables. Graph b) presents the most important variables, as measured by %IncMSE, highlighting the impact of factors such as vegetation cover and water availability on the accuracy of the model generated by Random Forest. These variables are not just important, they are fascinating in their impact on the Amazon’s tree density. Graph a) presents the most important variables measured by the IncNodePurity metric. The IncNodePurity metric refers to the increase in node purity (i.e., the ability of a node to separate classes) of a decision tree when a variable is used to split the data. The higher the IncNodePurity value, the greater the importance of the variable in the model. The results indicate that disturbance-related variables such as uspeed and lightning, as well as soil clay content, are the most influential in predicting the density of tall trees. This suggests that variations in disturbances and clay soils play a crucial role in the distribution of these trees, directly affecting their density. Graph b) illustrates the most important variables, as measured by %IncMSE. The %IncMSE metric represents the percentage increase in the Mean Squared Error (MSE) when a variable is excluded from the model. A higher value of %IncMSE indicates that the variable is crucial to the accuracy of the model. The most important variables according to this metric include climate factors such as mean annual precipitation and mean annual temperature. In addition, zonal wind speed (uspeed) and soil clay content indicate that they are key determinants in modeling the density of tall trees in the Amazon basin. These variables are not just data points but integral to our understanding of tree distribution in the Amazon.

**Figure S4:**
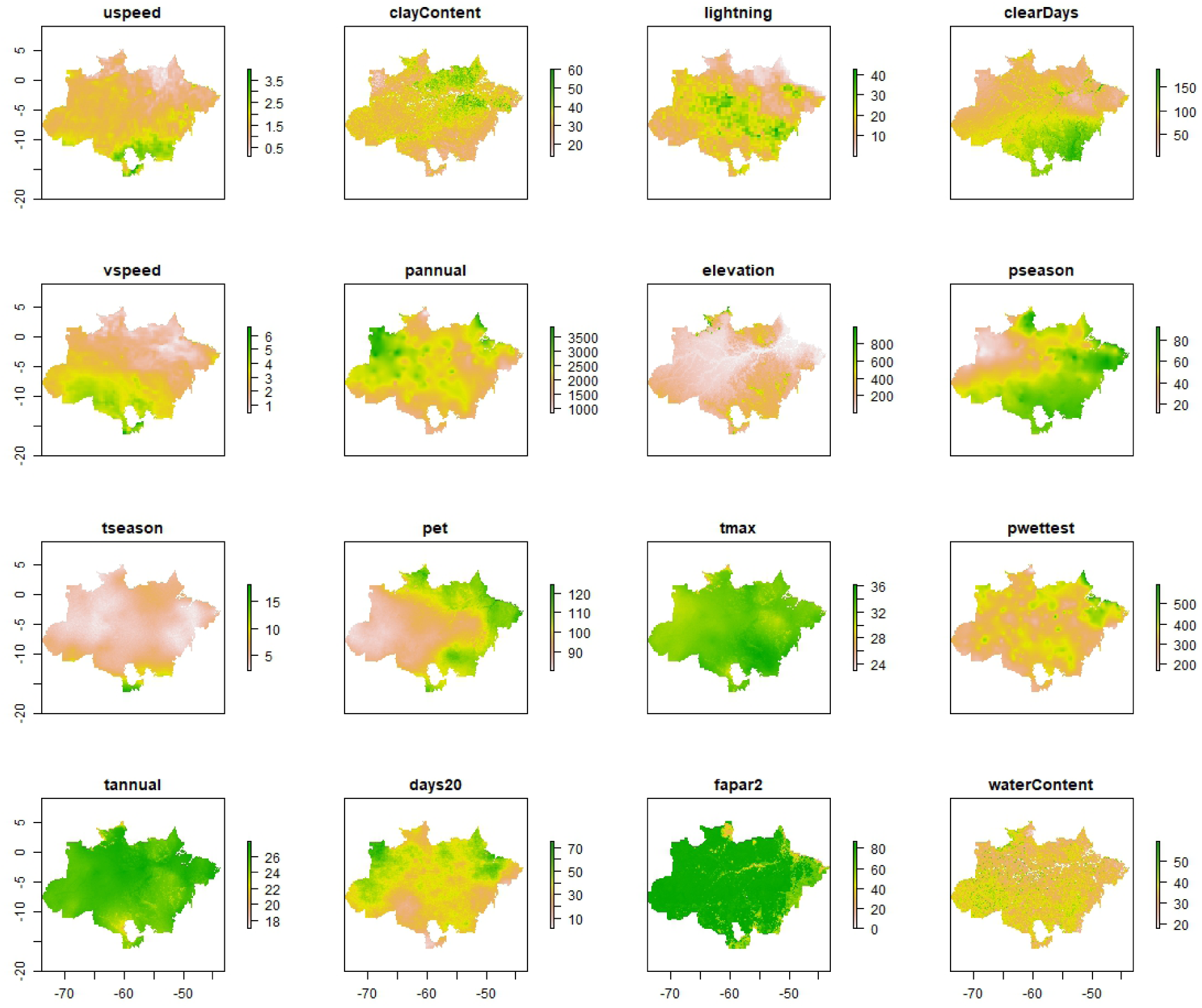
Maps of the 16 environmental variables used as predictors distributed in the Brazilian Amazon biome. Each panel represents the spatial distribution of a variable, highlighting its geographic differences. The mapped variables are uspeed and vspeed, wind speed components. clayContent: soil clay content. Lightning: lightning incidence. clearDays: number of clear days throughout the year. pannual: total annual precipitation. Elevation: terrain elevation. pseason and tseason: seasonality of precipitation and temperature, respectively. Pet: Potential evapotranspiration. Tmax: average annual maximum temperature. pwettest: precipitation in the wettest month. tannual: average annual temperature. days20: number of days with temperatures above 20°C. fapar2: fraction of photosynthetically active radiation absorbed by plants. waterContent: soil water content. The maps illustrate the spatial variation of each predictor, which is essential for environmental and ecological modeling in the Amazon region.

**Figure S5:**
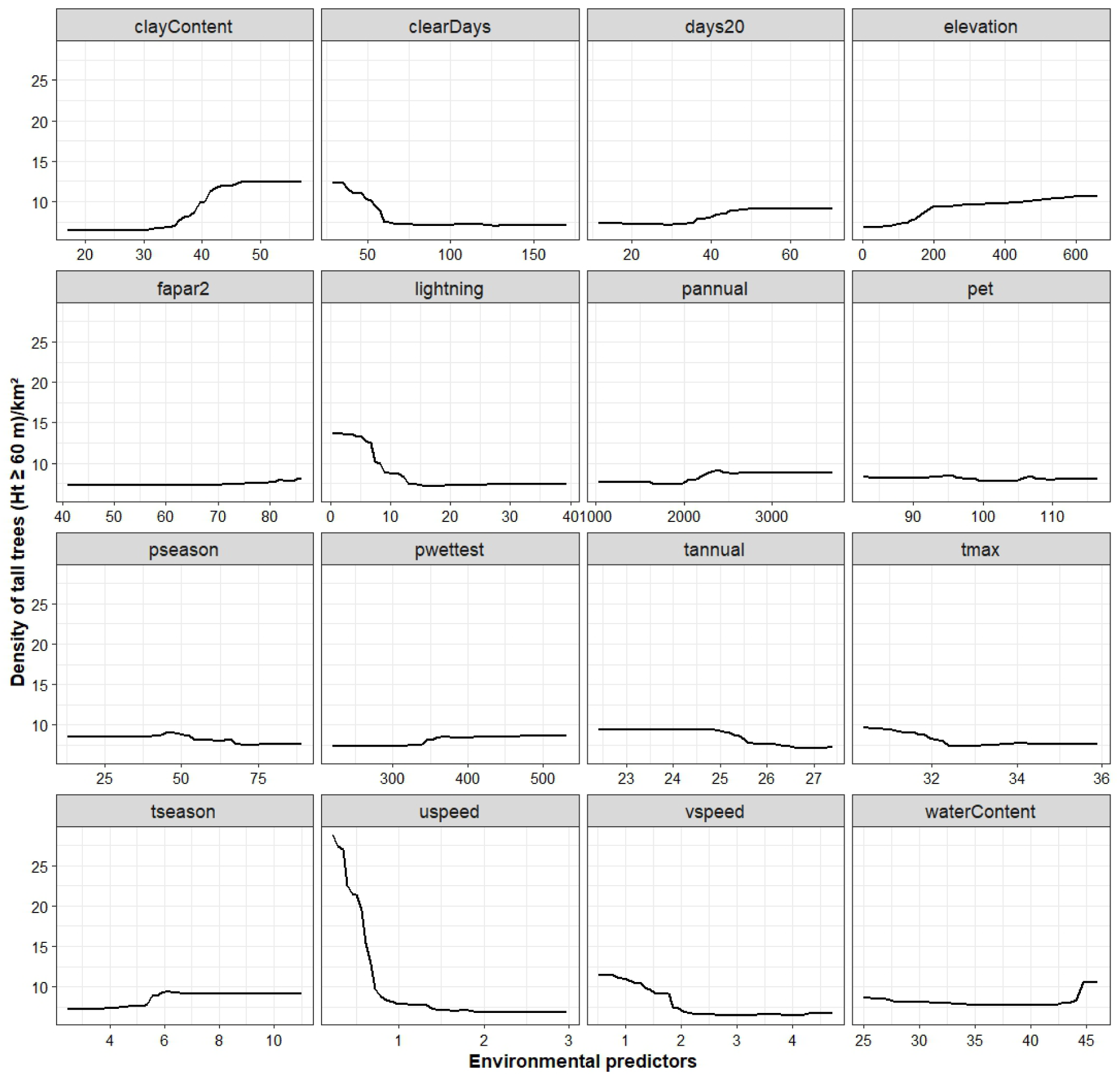
Partial dependence plot (pdp) for the environmental predictors of the Random Forest model for density of tall trees (⩾ 60 m) in the Amazon. In RF models, each predictor variable can interact with others in complex ways. The PDP is generated by calculating the average prediction of the model across all possible values of the target predictor, holding other variables fixed. For each value of the predictor (e.g., rainfall), the model computes the average predicted response (e.g., density of tall trees), providing a smoothed curve that shows how the response varies with changes in that predictor alone. This approach is particularly powerful in RF models because it helps mitigate the impact of multicollinearity, where predictor variables may be correlated with each other, complicating direct interpretations.

**Figure S6:**
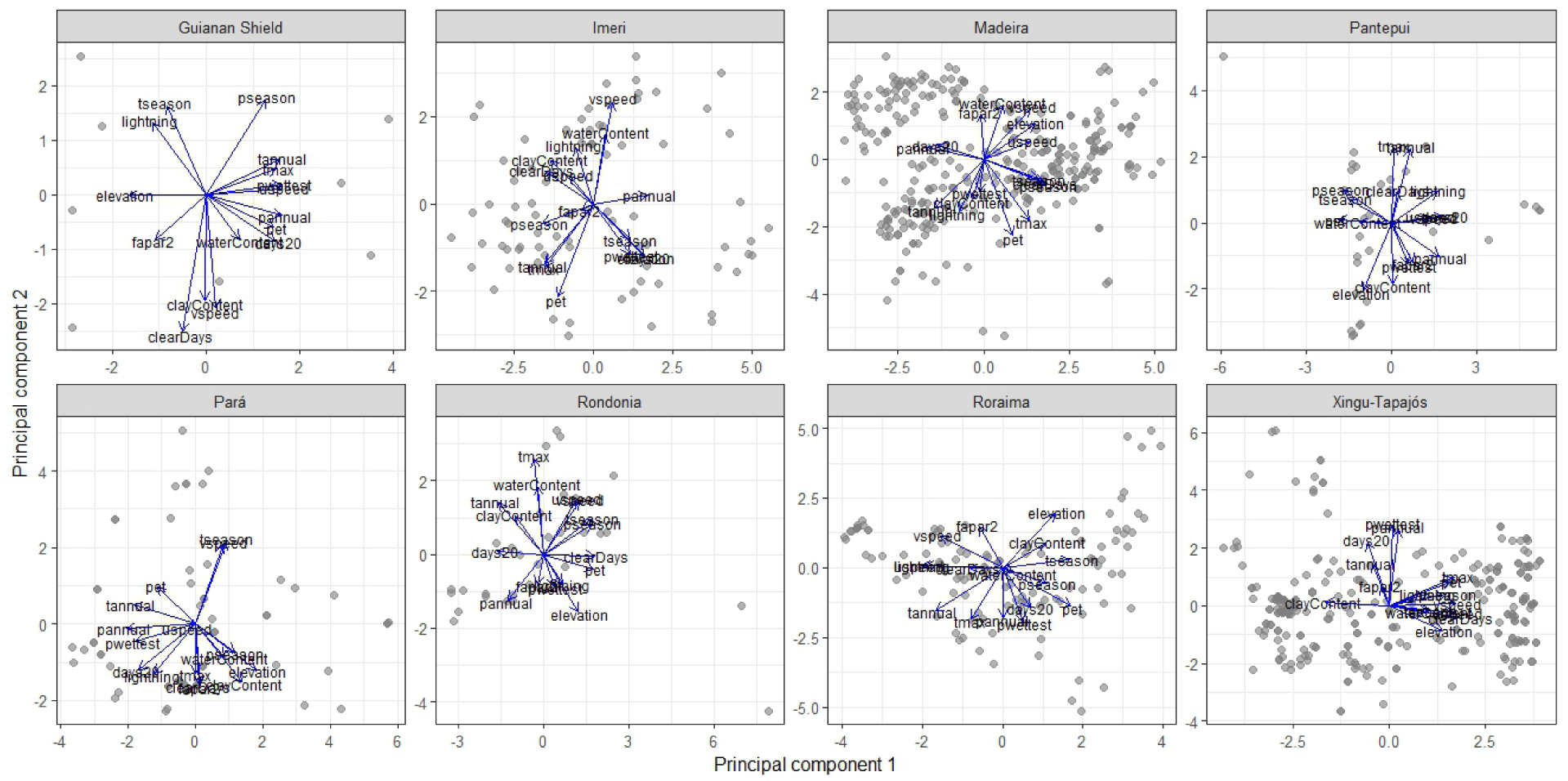
Principal components for environmental factors and biogeographic provinces: Guiana Shield (PC1 = 57.3%, PC2 = 17.6%), Pantepui (PC1 = 37.9%, PC2 = 25.9%) Roraima, (PC1 = 30.8%, PC2 = 19.8%), Imeri (PC1 = 41.4%, PC2 = 16.7%), Madeira (PC1 = 38.4%, PC2 = 15.8%), Rondonia (PC1 = 42.4%, PC2 = 17.1%), Xingu-Tapajós (PC1 = 42.3%, PC2 = 15.5%), Pará (PC1 = 34.3%, PC2 = 21.6%). Gray dots represent the data samples (containing the density per km2 and the underlying environmental factors). The position of each point along PC1 and PC2 indicates the combination of variables that define that point, that is, which geographic or environmental characteristics are dominant in that sample. We observe cluster points in a specific region of the graph, which may indicate that these samples share similar environmental characteristics. These clusters may reflect common geographic areas or environmental conditions. The arrows represent the environmental variables in the graph and indicate how these variables contribute to the principal components. Variables with longer arrows are more critical for the observed variation in the data. Variables close to each other or with aligned arrows are positively correlated. Variables (arrows pointing in opposite directions) are negatively correlated. The arrows’ length indicates the correlation’s strength with the principal components. Longer arrows indicate variables that contribute more to the variability along the corresponding principal component. If the arrow for one variable, for example, “elevation,” points to the right (positive PC1 axis) and another variable, such as “water content,” points to the top (positive PC2 axis), this means that these variables are vital contributors to explaining the variability in the data.

**Table S1:**
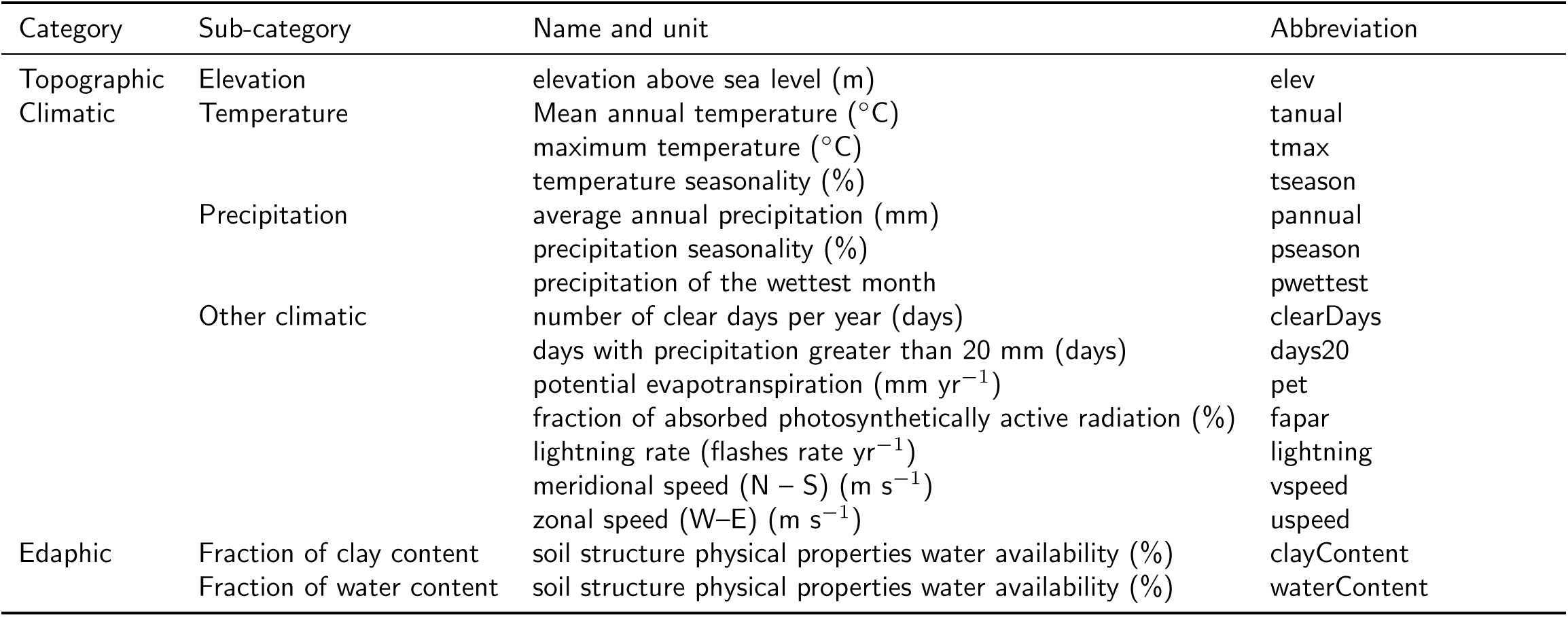
List of the main environmental variables selected for this study. Variable categories, subcategories, names and their corresponding units and abbreviations are shown.

